# Epidemiological survey of serum titers from adults against various Gram-negative bacterial V-antigens

**DOI:** 10.1101/718197

**Authors:** Mao Kinoshita, Masaru Shimizu, Koichi Akiyama, Hideya Kato, Kiyoshi Moriyama, Teiji Sawa

## Abstract

The V-antigen, a virulence-associated protein, was first identified in *Yersinia pestis* more than half a century ago. Since then, other V-antigen homologs and orthologs have been discovered and are now considered vital molecules for the toxic effects mediated by the type III secretion system in infections caused by various pathogenic Gram-negative bacteria. After purifying recombinant V-antigen proteins including PcrV from *Pseudomonas aeruginosa*, LcrV from *Yersinia*, LssV from *Photorhabdus luminescens*, AcrV from *Aeromonas salmonicida*, and VcrV from *Vibrio parahaemolyticus*, we developed an enzyme-linked immunoabsorbent assay to measure titers against each V-antigen in the sera collected from 186 adult volunteers. Different titer-specific correlation levels were determined for the five V-antigens. The anti-LcrV and anti-AcrV titers shared the highest correlation with each other, with a correlation coefficient of 0.84. The next highest correlation coefficient was between anti-AcrV and anti-VcrV titers at 0.79, while the lowest correlation was found between anti-LcrV and anti-VcrV titers, which were still higher than 0.7 Sera from mice immunized with one of the five recombinant V-antigens displayed cross-antigenicity with some of the other four V-antigens, supporting the results from the human sera. Thus, the serum anti-V-antigen titer measurement system could potentially be used for epidemiological investigations on various pathogenic Gram-negative bacteria.

## Introduction

The type III secretion system (TTSS) plays an important role in the virulence of many Gram-negative bacteria [1-3]. Through the TTSS, Gram-negative bacteria inject their effector molecules to target eukaryotic cells and induce a favorable environment for their infections. Translocation is a mechanism through which effector molecules of the TTSS pass through the targeted eukaryotic cell membrane. During translocation, three bacterial proteins form a translocational pore structure called the ‘translocon’ [1-3]. A cap protein in the secretion apparatus of the type III secretion needle is one type of translocon protein, which for historical reasons is called the V-antigen in *Yersinia* spp. [4-8]. Briefly, the V-antigen, a virulence-associated protein, was identified as an antigenic component recognized by the immune system in *Yersinia pestis* plague-infected hosts more than half a century ago [4-8]. In the 1980s, the V-antigen of *Y. pestis* was identified as a low-calcium response (*lcr*)-associated protein (named LcrV) encoded in the plasmid associated with its virulence [9]. A homologous gene called PcrV was identified in the *Pseudomonas aeruginosa* genome [10], and it has been reported that the virulence associated with the TTSS can be inhibited by a specific antibody against LcrV in *Yersinia* and PcrV in *P. aeruginosa* [11, 12]. Because vaccinating mice with LcrV or PcrV had protective effects against lethal infections with *Yersinia* or *P. aeruginosa*, respectively, later on, anti-PcrV immunotherapy was developed to target human infections with *P. aeruginosa* using immunoglobulins [13-24] and vaccines [25-27], from which and several projects have progressed to human clinical trials [28-31].

We recently published an epidemiological study on serum titers against PcrV in human volunteers [32], and another showing how the prophylactic administration of human serum-derived immunoglobulin with a high anti-PcrV titer significantly improved the survival rate, pulmonary edema, and inflammatory cytokine production in a *P. aeruginosa* pneumonia model [18]. The results of both studies imply that immunity against the V-antigen and its homologs might be necessary for preventing infections caused by pathogenic bacterial species utilizing the TTSS-virulence mechanism. The possession of V-antigen homologs has recently been reported in several Gram-negative bacteria, including *Aeromonas* spp., *Vibrio* spp., and *Photorhabdus luminescens* (hereafter referred to as *Ph. luminescens*) [33]. Although specific immunity against the V-antigen or its homologs seems to be important for host immunity against such bacterial infections, insufficient information is available on human immunity against V-antigens. Therefore, here, we conducted epidemiological studies on serum titers against the V antigen and its homologs in *Y. pestis, Aeromonas salmonicida, Vibrio parahaemolyticus*, and *Ph. luminescens.* Potential associations for age, titer levels, and cross-reactivity among the recombinant V-antigen homologs were evaluated, and for some species the titer levels against these antigens were highly correlated and some V-antigen homologs showed cross-reactivity. Speculation about specific immunity against multiple pathogens will also be discussed.

## Materials and methods

### Construction of five recombinant Gram-negative bacterial V-antigens (PcrV, LcrV, AcrV, VcrV, and LssV)

Five recombinant V-antigens were constructed. Details on the PCR primers and cloning sites are listed in **Table 1**. The coding regions in the V-antigens were polymerase chain reaction (PCR)-amplified with specific primers containing the restriction enzyme sites for insertion into a protein expression vector. PCR-amplified genes were cloned into the pCR2.1 cloning vector and *E. coli* TOP10F cells via TOPO cloning (Thermo Fisher Scientific, Waltham, MA, USA). After digesting the purified plasmids containing each individual cloned gene with restriction enzymes, the inserted coding regions of each gene were transferred to the multi-cloning site of expression vector pQE30 (Qiagen, Hilden, Germany) for the expression of a hexahistidine-tagged protein in *E. coli* M15. The various endotoxin-free Gram-negative bacteria V-antigens were prepared as reported previously (**Fig. 1**) [17, 20].

**Table 1.**
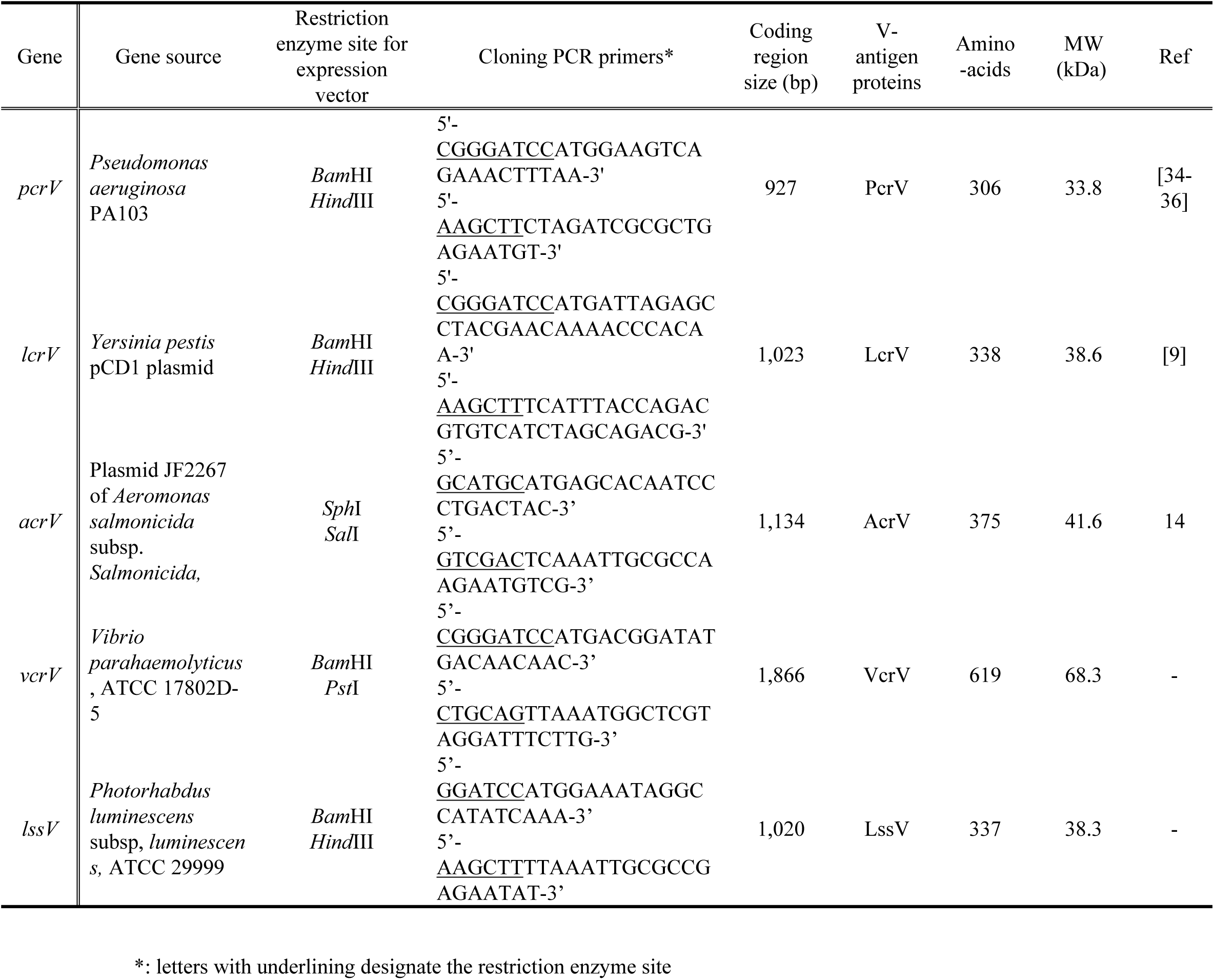
Gene sources, primer sets for V-antigen gene cloning, and characteristics of the recombinant V-antigens used in this study.

**Fig. 1.**
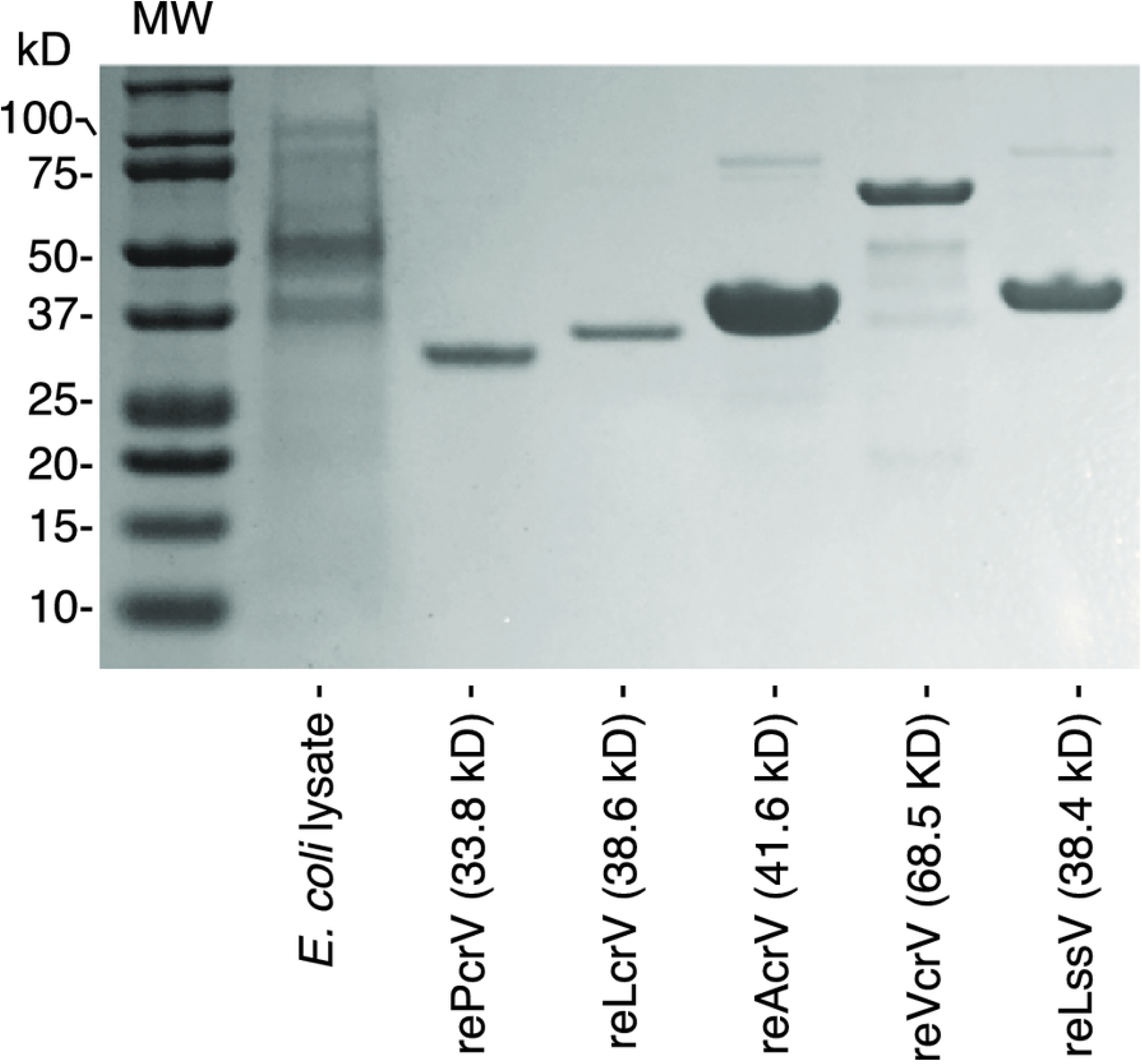
Sodium dodecyl sulfate-polyacrylamide gel electrophoresis (SDS-PAGE) separation of extracted recombinant hexahistidine-tagged V-antigen proteins. Recombinant PcrV from *P. aeruginosa*, LcrV from *Y. pestis*, AcrV from *Aeromonas salmonicida*, VcrV from *V. parahaemolyticus*, and LssV from *Ph. luminescens* were SDS-PAGE separated using a 10% Bis-Tris-gel.

### Survey participants and study background

This study was approved by the Kyoto Prefectural University of the Medicine Ethics Committee. Surgical adult patients (n=186) who were undergoing anesthesia in the central operating division of the Kyoto Prefectural University of Medicine, from April 2012 to March 2013, participated in this study as volunteers, as previously reported [32]. Briefly, serum was prepared from the remaining small amount of each blood sample collected for arterial blood gas analysis on the induction of anesthesia, and then stored at −80°C.

### Anti-V antigen titer measurements

We developed an enzyme-linked immunoabsorbent assay (ELISA) for measuring the serum anti-V antigen titers. Micro-well plates (Nunc C96 Maxisorp; Thermo Fisher Scientific) were coated for 2 h at 4°C with recombinant V-antigen proteins (recombinant PcrV from *P. aeruginosa*, LcrV from *Yersinia*, AcrV from *Aeromonas*, VcrV from *Vibrio*, and LssV from *Ph. luminescens*) suspended in coating buffer (1.0 μg/mL in coating solution; 0.0M NaHCO_3_, pH 9.6) [32] (**Fig.1**). The plates were washed twice with phosphate-buffered saline (PBS) containing 0.05% Tween-20 (P9416; Sigma-Aldrich, St. Louis, MO, USA), and then blocked with 200 mL of 1% bovine serum albumin/PBS overnight at 4°C. Samples (serial dilution: 1280×) were applied to the plates (100 mL/well) and then incubated overnight at 4°C. Peroxidase-labeled anti-human IgG (A8667, Sigma-Aldrich) was applied at 1:60,000 for 1 h at 37°C. After six washes, the plates were incubated with 2,2’-azino-bis (3-ethylbenzthiazoline-6-sulfonic acid) (A3219; Sigma-Aldrich) at room temperature for 30 min. After adding 0.5 M H_2_SO_4_ at 100 mL/well to the plates, the optical density (O.D) at 450 nm was measured with a microplate reader (MTP-880Lab; Corona Electric, Hitachinaka, Japan). To ensure no cross-reactivity with *Escherichia coli* proteins, competitive ELISA with a soluble fraction of *E. coli* M15 lysate was performed, and no significant effect on titer measurement was observed.

### Immunizing mice with the V-antigens

Certified pathogen-free, male ICR mice (4 weeks old) were purchased from Shimizu Laboratory Supplies, Co, Ltd., Japan. Mice were housed in cages with filter tops under pathogen-free conditions. The protocols for all animal experiments were approved by the Animal Research Committee of Kyoto Prefectural University of Medicine, Kyoto, Japan, before undertaking the experiments (authorization number: M29-592). Three mice per group were intradermally immunized with one of the five recombinant V-antigen proteins (10µg/dose) adjuvanted with complete Freund’s adjuvant in the first injection, and four weeks later with incomplete Freund’s adjuvant as a second injection. Eight weeks after the first injection, the immunized mice were euthanized, and peripheral blood samples were collected from them. Serum titers against the five V-antigens were individually measured by ELISA, as described above.

### Phylogenetic and cluster analyses

The five V-antigens were phylogenetic analyzed using ClustalW (Genome Net, https://www.genome.jp/tools-bin/clustalw), or RStudio (version 1.2, Boston, MA, USA. https://www.rstudio.com) with R version 3.4.3 (The R Foundation, https://www.r-project.org). Unrooted trees were prepared using the neighbor-joining method and rooted trees were prepared using the unweighted pair group method with arithmetic mean, and heat maps where the individual values contained in a matrix are represented as colors with dendrograms were made using the heat map function of R language. The predicted three-dimensional structures were generated with the Cn3D macromolecular structure viewer at the National Center for Biotechnology Information (https://www.ncbi.nlm.nih.gov/Structure/CN3D/cn3d.shtml).

## Results

### Anti-V antigen titers and volunteer age distribution

The 186 volunteers consisted of 111 (59.7%) males and 75 (40.3%) females, and the following age distribution: 20–29 years (13, 7.0%), 30–39 years (20, 10.8%), 40–49 years (26, 14.0%), 50–59 years (23, 12.4%), 60–69 years (40, 21.5%), 70–79 years (42, 22.5%), and 80 years plus (22, 11.8%). No study participant had an active infection. The titers against the V-antigen across the age distribution of the 186 participants are shown in **Fig. 2.** There was no statistically significant correlation in the linear regression between age and each anti-V antigen titer, although high anti-PcrV titers were more common at over 50 years of age in the population, as reported previously [32]. As an overall trend, two separate titer peaks in the 40–50 and 70–80 age groups were observed in the age distribution of all the anti-V-antigen titers.

**Fig. 2.**
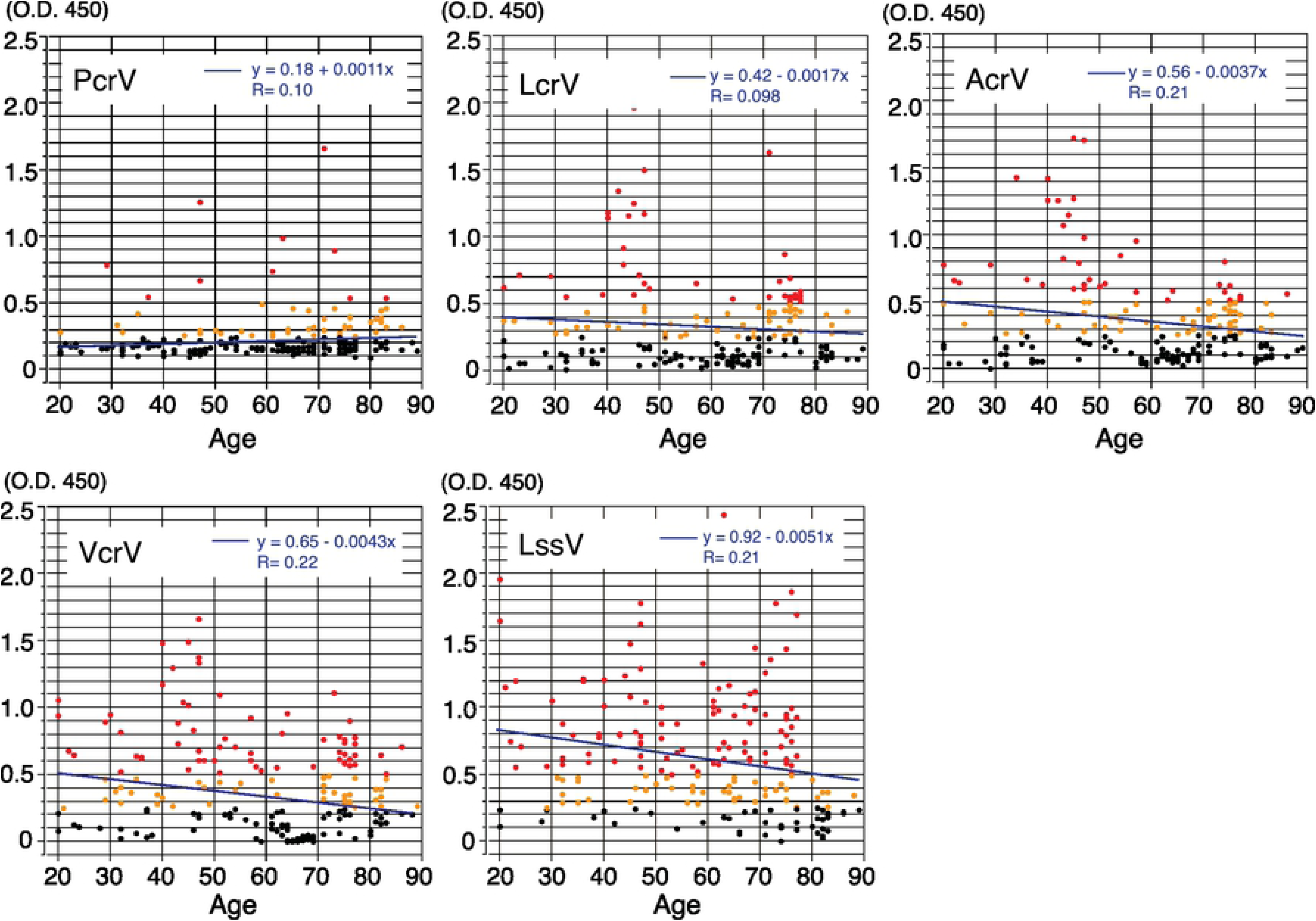
Age distribution of human V-antigen titers. The diluted serum (1,280×) was used for ELISAs, and the O.D. 450 nm values were measured. O.D. values of 0.5 or more are indicated by red dots, while yellow dots represent values between 0 and 0.3.

### Correlations among the anti-V antigen titers

We next analyzed whether any correlations existed among the five anti-V antigen titers. The anti-LcrV and the anti-AcrV titers showed the highest correlation with a correlation coefficient 0.84, followed by the anti-AcrV and anti-VcrV titers at 0.79, and the anti-LcrV titers and anti-VcrV titers at 0.74 (**Fig. 3**) No statistically significant correlations were detected between anti-PcrV and any of the other anti-V-antigens.

**Fig. 3.**
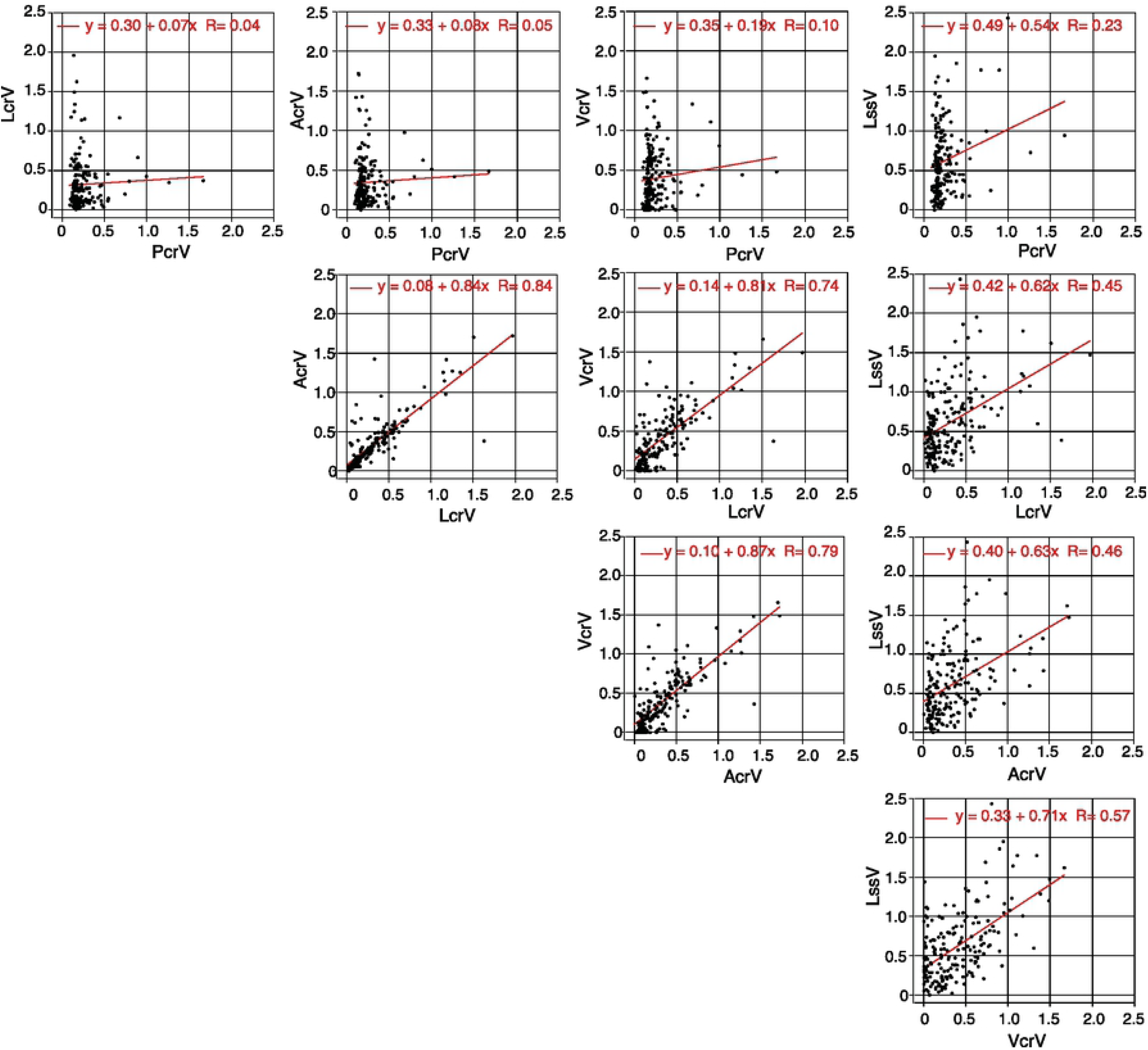
Serum titer correlations between two V-antigens. The diluted serum (1,280×) was used for ELISAs, and the O.D. values at 450 nm were measured. The titer correlation of the two V-antigens was mapped in an X–Y plot.

A cluster analysis was prepared from the correlation coefficient values, and phylogenetic trees and a heat map were made (**Fig 4A**). The unrooted and rooted phylogenic trees and the heat map show that anti-PcrV had the most unique profile among the five anti-V-antigen titers. The anti-LssV titer is located between anti-PcrV and three other anti-V-antigen titers. Higher homology in antigenicity was observed among anti-AcrV, anti-LcrV, and anti-VcrV titers.

**Fig. 4.**
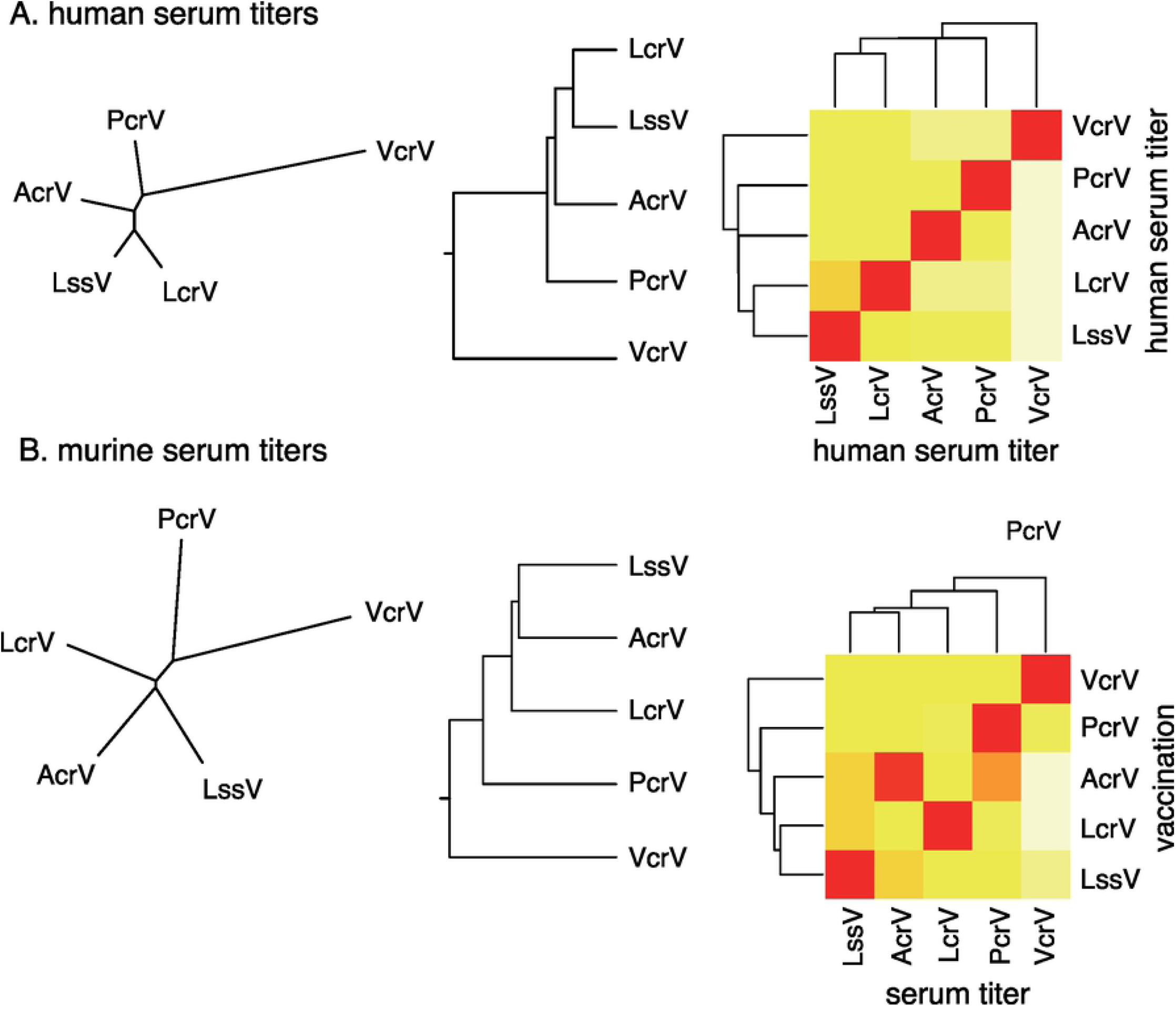
Phylogenetic trees and heat maps showing the correlations among the five anti-V-antigen titers. **A.** Human serum titers. The correlation coefficients of the serum titer correlations shown in **Fig. 3** were matrixed. Phylogenetic trees (an unrooted tree, neighbor-joining method; and a rooted tree, unweighted pair group method with arithmetic mean) and a heat map were constructed. **B.** Anti-V-antigen serum titers from mice immunized with one of the five V-antigens. Phylogenetic trees (unrooted tree, neighbor-joining method; and a rooted tree, unweighted pair group method with arithmetic mean) and a heat map were constructed.

### Cross-reactive antibodies against V-antigens and anti-V antigen titers in the serum from immunized mice

To evaluate potential cross-antigenicity among the five V-antigens, the serum titers from a mouse immunized with one of five recombinant V-antigen proteins were measured by ELISA (**Fig 5**). Cluster analysis was applied to the analysis by using the O.D. values from the ELISA, from which phylogenetic trees and a heat map were made (**Fig 4B**). LssV and AcrV showed higher cross-antigenicity with each other, unlike VcrV, which was less cross-reactive against the other V-antigens (**Fig. 4B and 5)**. While the sera from the AcrV-immunized mice reacted with PcrV, the sera from PcrV-immunized mice did not react with AcrV.

**Fig. 5.**
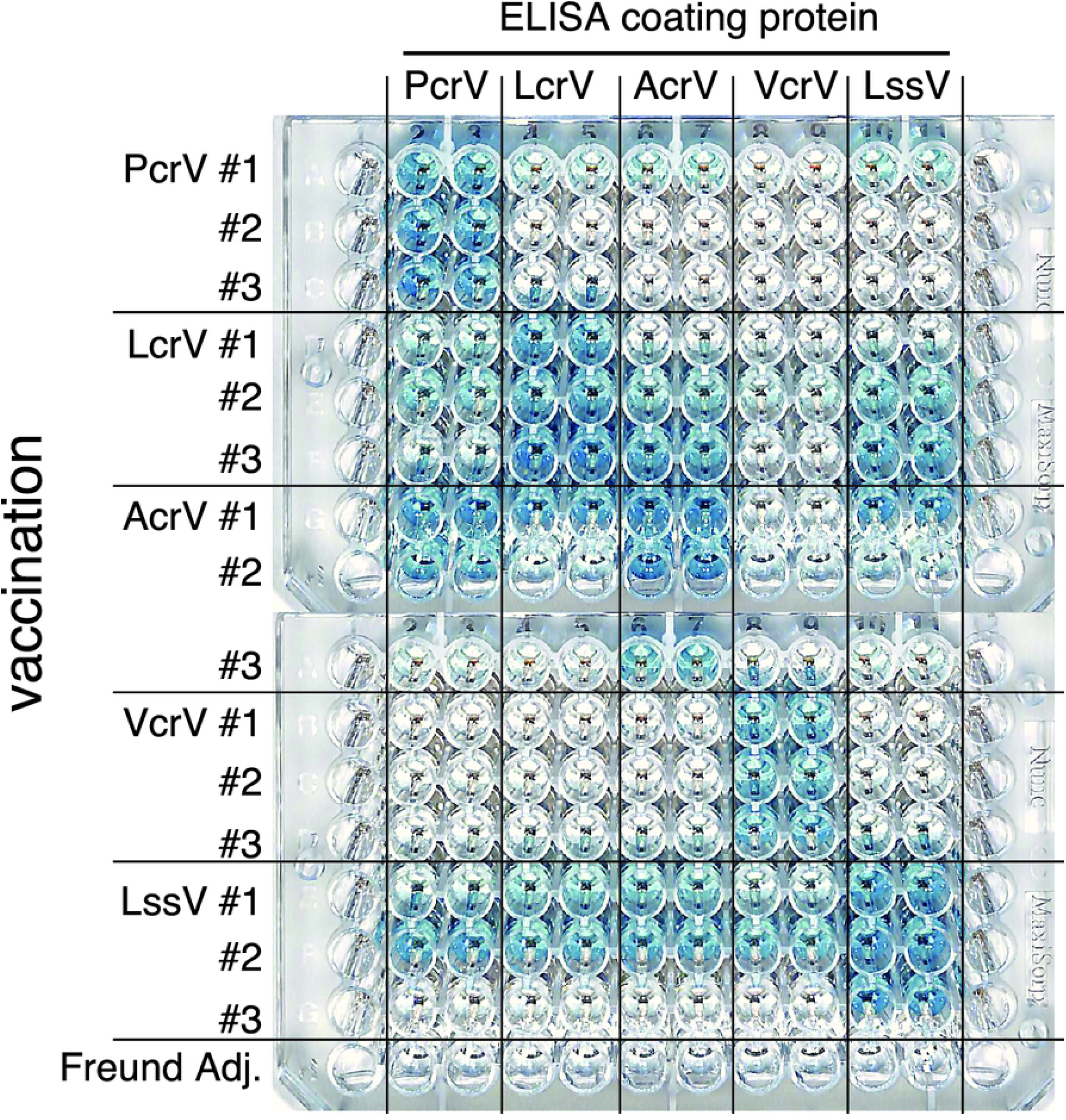
ELISA plate displaying the anti-V-antigen serum titers from mice immunized with one of the five V-antigens. Three mice per group were intradermally immunized with one of five recombinant V-antigen proteins (10µg/dose) adjuvanted with complete Freund’s adjuvant as the first immunization and, four weeks later, with incomplete Freund’s adjuvant at as the second immunization. Eight weeks after the first immunization, mice were euthanized and peripheral blood samples were collected. Serum titers against the five V-antigens were measured by ELISA. The plates were incubated with 2,2’-azino-bis (3-ethylbenzthiazoline-6-sulfonic acid) at room temperature for 30 min, and this photo was taken.

Next, to determine whether correlations among the V-antigens have some association with the identity of the primary protein sequences, we investigated the identity of these sequences for the five V-antigens using the BLOSUM substitution score matrix in ClustalW (**Fig. 6**). In this map, the overall similarities of two out of five of the V-antigens for the whole molecules ranged between 21 and 49%. The similarity of the amino-terminal domain (14–51%) and central domains (14–45%) can be seen to be low in comparison with the carboxyl-terminal domain that showed 48–84% similarity. VcrV is unique with a long extra sequence (over 160 amino acids, aa) at the amino-terminal domain, and an extra sequence (of 80 aa) in the central domain of the sequence alignment.

**Fig. 6.**
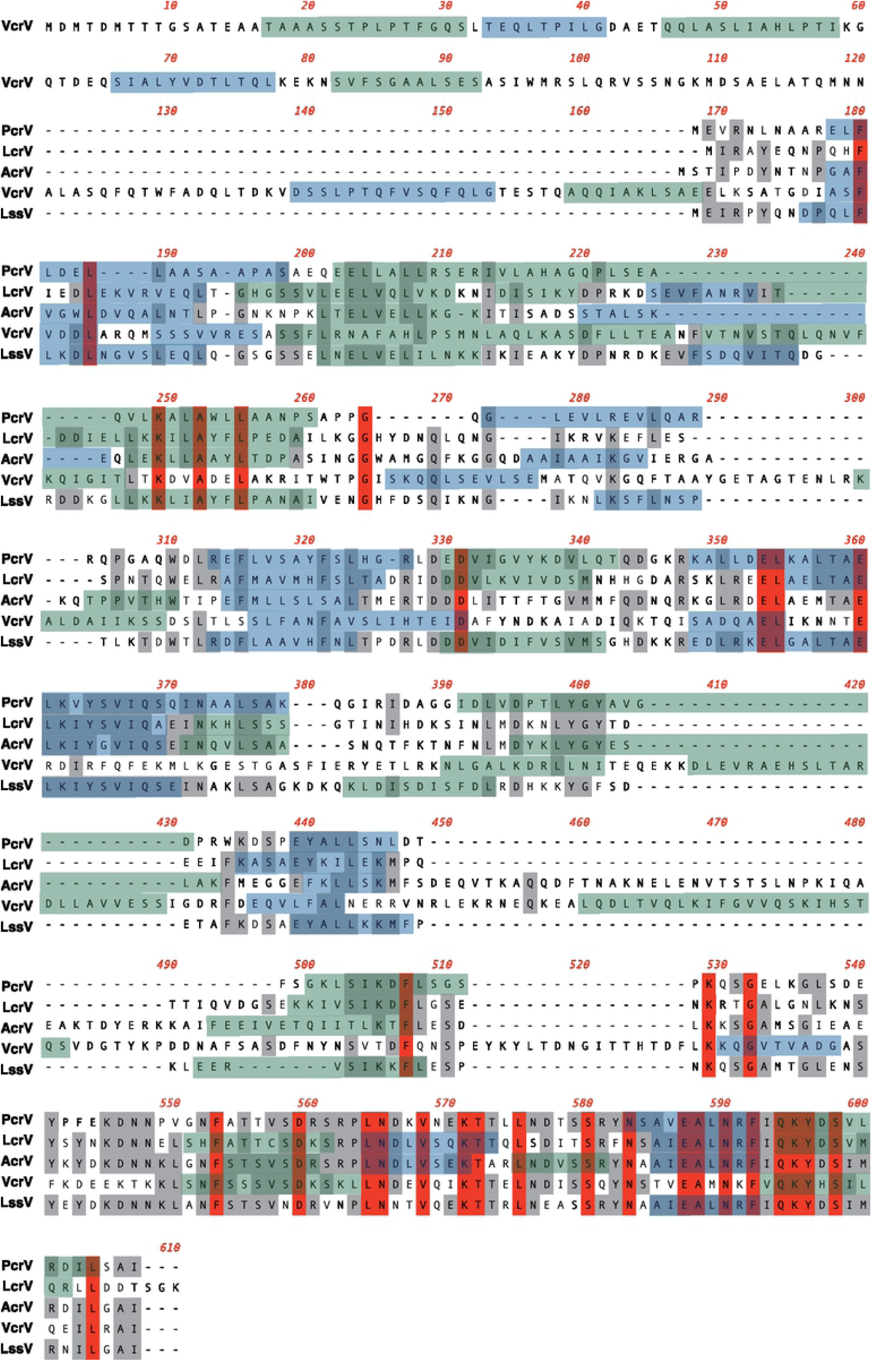
Clustal W sequence alignment of the five primary V-antigen sequences. The overall sequence similarities of two of the five V-antigens across the whole molecules are between 21 to 49%. The similarity of the amino-terminal domain (14–51%) and central domains (14–45%) is low in comparison with the carboxyl-terminal domain, which has a 48–84% similarity range. VcrV is unique with an extra-long amino-terminal domain sequence (over 160 aa), and an extra sequence (80 aa-) in the central domain of the sequence alignment.

We also made phylogenetic trees of V-antigens based on the primary amino acid sequence similarity scores of whole molecules, amino-terminal domains, central domains, and carboxyl-terminal domains (**Fig. 7**). In the cluster analysis of the carboxyl domains with the highest similarity scores (48–84%) for their amino-acid sequences, AcrV, LssV, and PcrV are closer than VcrV and LcrV. These phylogenetic analyses using the primary amino-acid sequences did not match well with the phylogenetic trees constructed from the correlations among the serum IgG titers of the five V-antigens (**Fig. 4A**). In particular, VcrV, which has an extremely long central domain containing regions that are missing in the other V-antigens, occupies a separate position in the phylogenetic trees made from the primary amino-acid sequences. Therefore, these findings suggest the titer cross-antigenicity in human sera may not be correlated with similarity in the amino-acid primary sequences.

**Fig. 7.**
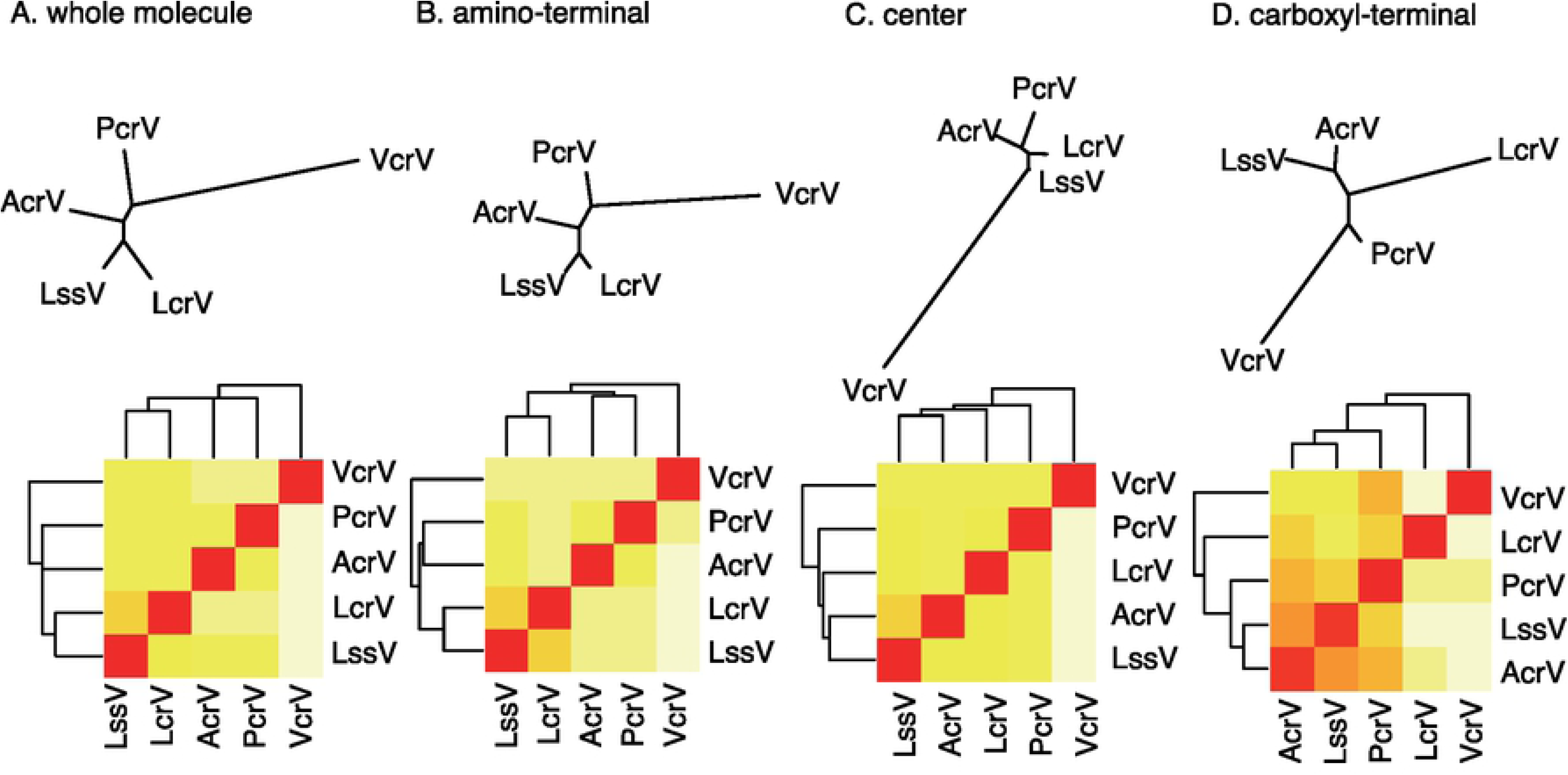
Phylogenetic trees and heat maps based on the primary amino-acid sequences (whole molecule, amino-terminal, center, and carboxyl-terminal) of the V-antigens. Phylogenetic analyses of the primary amino-acid sequences of whole molecules, amino-terminal, center, and carboxyl-terminal domains. A. Complete primary sequence. B. Amino-terminal domain, C. Center domain, D. Carboxyl-terminal domain. The amino acid positions in the amino-terminal, center, and carboxyl-terminal domains are as follows: LcrV: #1-#164, #165-#278, #279-#326 respectively; PcrV:#1-#142, #143-#256, #257-#294, respectively; AcrV: #1-#162, #163-#316, #317-#361, respectively; VcrV:#1-#361, #362-#558, #559-#607, respectively; LssV: #1-#170, #171-#280, #281-#325, respectively.

As we have previously reported, a blocking monoclonal antibody (called Mab166) against PcrV recognizes a conformational structure but not a primary amino-acid sequence [15]. As shown in **Fig. 8**, the predicted three-dimensional structures of the V-antigens have similar dumbbell-like structures with two globular domains on either end of a grip formed by two coiled-coil motifs [37]. The grip that connects the two globular domains contains an antiparallel coiled-coil structure comprising a central coiled-coil region and a carboxyl-terminal coiled-coil region. The carboxyl-terminal is folded into a single long α-helix. Therefore, there is a possibility that the correlations we have observed may be dependent on the sequence similarity of the three-dimensional structures associated with the conformational epitopes.

**Fig. 8.**
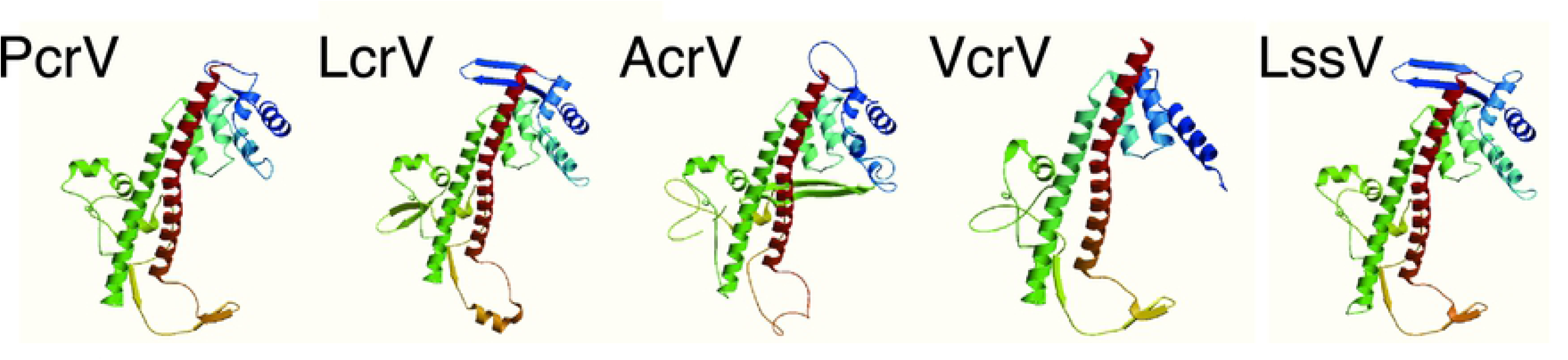
The predicted three-dimensional V-antigen structures. The predicted three-dimensional structures were generated by the Cn3D macromolecular structure viewer at the National Center for Biotechnology Information (https://www.ncbi.nlm.nih.gov/Structure/CN3D/cn3d.shtml).

## Discussion

*Yersinia* LcrV has been recognized as a V-antigen with immunoprotective characteristics in *Yersinia* infections since the 1950s [4, 5, 7, 8], but it took almost 50 years until LcrV, *P. aeruginosa* PcrV, and *Aeromonas* AcrV were anatomically visualized as distinct structures on the tip of the needle of the injectisome of the type III secretion apparatus [38-41]. Because specific antibodies binding to a particular portion of this structure can inhibit the translocation of type III secretory toxins in *Yersinia* and *P. aeruginosa* [11, 42], gaining better understanding of the interactions between V-antigens and the host’s humoral immunity against the virulence of bacterial type III secretion is important if potential non-antibiotic treatments for infections in various hosts are to be developed.

Cross-antigenicity among *Yersinia* spp. was reported nearly 40 years ago. In 1980, cross-immunity to *Y. pestis* was noted in mice that had been orally infected with *Y. enterocolitica* serotype O3 [43]. In 1983, it was also reported that partial protection in mice against *Y. pestis* infection by the *Y. enterocolitica* V-antigen was linked to the partial cross-reactivity of V-antigens [44]. Later, DNA sequencing of the most common serotypes of human pathogenic *Y. enterocolitica* and *Y. pseudotuberculosis* revealed that two evolutionary distinct types of V-antigen exist in *Yersinia* spp. [45]. One type is represented by the *Y. enterocolitica* serotype O8 strains WA, WA-314, and NCTC 10938 (LcrV-YenO8), while the other type comprises *Y. pestis, Y. pseudotuberculosis*, and *Y. enterocolitica* serotypes O3, O9, and O5 (LcrV-Yps). By raising monospecific antisera against both types of V-antigen (*Y. enterocolitica* serotypes O3 and O8), anti-V-antigen serum was protective only if the immunizing V-antigen was the same type as the V-antigen produced by the infecting strain. The difference in protective immunity between the two types was caused by the presence of a hypervariable region between amino acids 225 and 232. The protectivity of the V-antigen was later confirmed and refined using the recombinant V-antigen of *Y. pseudotuberculosis* and monospecific anti-V-antigen serum [46]. In that study, antiserum against the *Y. pseudotuberculosis* V-antigen provided mice with passive immunity to a challenge infection with *Y. pestis* or *Y. pseudotuberculosis* but not *Y. enterocolitica* O8 (strain WA). These past studies on cross-antigenicity to *Yersinia* V-antigens in mice imply that the structure of the key domain, but not the overall primary amino acid sequence similarity *per* se, is important for protective immunity.

Other than *Yersinia* LcrV, only two studies about *P. aeruginosa* PcrV have reported on antibody titers against *P. aeruginosa* PcrV in human serum [32, 47]. No other studies investigating antibody titers against other V-antigens have been reported as yet. *V. parahaemolyticus*, a Gram-negative marine bacterium, the V-antigen titers against it of which were examined in the present study, causes food-borne gastroenteritis [48, 49]. Among *Vibrio* spp., *V. harveyi* (a Gram-negative bioluminescent marine bacterium) is ubiquitous in the marine environment. It is considered one of the important bacterial species that form the normal flora of healthy shrimp and it carries a V-antigen homolog gene in its genome [50]. This bacterium is sometimes recognized as causing high mortality in shrimps in the worldwide shrimp fishing industry [51, 52]. As a halophilic *Vibrio* species, *V. alginolyticus*, which causes wound infections, was first recognized as pathogenic to humans in 1973 [53]. Recent studies have proposed that *V. alginolyticus* possesses the same TTSS gene organization as *V. parahaemolyticus* and *V. harveyi* [54, 55]. *Aeromonas* spp., such as *A. salmonicida* and *A. hydrophila*, are not pathogenic to humans but cause infections in salmon and trout and carry the AcrV V-antigen homolog. However, *A. hydrophila* sometimes causes gastroenteritis in humans, who acquire infections with it by ingesting food or water containing sufficient numbers of this organism [56-58]. *Ph. Luminescens* (a gammaproteobacterium within the *Enterobacteriaceae* family) is also not pathogenic to humans, but is a lethal pathogen of insects and possesses a pathogenicity island encoding the TTSS including the LssV V-antigen homolog [59-62]. It lives in the gut of an entomopathogenic nematode within the *Heterorhabditidae* family. However, human infections with *Photorhabdus* spp. have recently been reported in the USA and Australia, suggesting that these bacteria are emerging human pathogens [63]. In comparison with the infections caused by *P. aeruginosa*, infections caused by *Y. pestis* or *V. parahaemolyticus* are less common. This makes it interesting that titers against V-antigens from non-human pathogens were also detected in this study. Anti-AcrV showed a high correlation with anti-LcrV, and some correlation was detected between anti-LssV and anti-VcrV. The titers against the antigens from non-human pathogens might result from cross-antigenicity among the V-antigen homologs. Our cluster analysis with a heat map of anti-V-antigen titers from the 182 adult volunteers displayed a higher correlation between LcrV, AcrV, and VcrV than the correlation between PcrV and LssV and the other V-antigens (**Fig. 4A**).

Immunizing mice with one of the five recombinant V-antigens resulted in cross-antigenicity of the sera against the V-antigens (**Figs. 4B** and **5)**. This result shows that VcrV is unique, while AcrV, LssV, and LcrV share some degree of cross-antigenicity with each other, although the results differed slightly from the cross antigenicity observed with human sera. In our previous experiments on anti-PcrV titers in human sera, the administration of extracted IgG derived from high titer anti-PcrV sera protected against challenge with lethal pneumonia in a murine model [18]. Therefore, despite no similar immunological experiments being performed with other V-antigens, our study has shown that the cross-reactive antigenicity we observed among the various V-antigens with serum-specific anti-V antigen titers could potentially afford some degree of immunological protection against various Gram-negative bacteria. Further investigation of the conformational blocking epitopes in the needle cap structure of the type III secretory apparatus and the immunological aspects of the structural antigenicity of a critical portion of the V-antigens should provide better understanding on how to effectively block the TTSS-associated virulence associated with various Gram-negative pathogens.

## Acknowledgments

This work was supported by grants from Grant-in-Aid for Scientific Research (KAKENHI No. 24390403, 26670791, 23659748, and 20816384) and the Ministry of Education, Culture, Sports, Science and Technology of Japan to Teiji Sawa. We thank Sandra Cheesman, PhD, from Edanz Group (www.edanzediting.com/ac) for editing a draft of this manuscript.

## Author Contributions

**Conceptualization:** Teiji Sawa.

**Data curation:** Mao Kinoshita, Teiji Sawa.

**Formal analysis:** Mao Kinoshita, Teiji Sawa.

**Funding acquisition:** Teiji Sawa.

**Investigation:** Mao Kinoshita, Teiji Sawa.

**Methodology:** Mao Kinoshita, Masaru Shimizu, Koichi Akiyama, Hideya Kato.

**Project administration:** Teiji Sawa.

**Resources:** Masaru Shimizu, Kiyoshi Moriyama.

**Supervision:** Teiji Sawa.

**Writing – original draft:** Mao Kinoshita, Teiji Sawa.

**Writing – review & editing:** Kiyoshi Moriyama, Teiji Sawa.

## References

1. Cornelis GR. The type III secretion injectisome, a complex nanomachine for intracellular ‘toxin’ delivery. Biol Chem. 2010;391(7):745–51. doi: 10.1515/BC.2010.079 PMID: 20482311

2. Erhardt M, Namba K, Hughes KT. Bacterial nanomachines: the flagellum and type III injectisome. Cold Spring Harb Perspect Biol. 2010;2(11):a000299. doi: 10.1101/cshperspect.a000299 PMID: 20926516

3. Galan JE, Lara-Tejero M, Marlovits TC, Wagner S. Bacterial type III secretion systems: specialized nanomachines for protein delivery into target cells. Annu Rev Microbiol. 2014;68:415–38. doi: 10.1146/annurev-micro-092412-155725 PMID: 25002086

4. Bacon GA, Burrows TW. The basis of virulence in *Pasteurella pestis*: an antigen determining virulence. Br J Exp Pathol. 1956;37(5):481–93 PMID: 13374206

5. Burrows TW. An antigen determining virulence in *Pasteurella pestis*. Nature. 1956;177(4505):426–7 PMID: 13309325

6. Burrows TW. Biochemical properties of virulent and avirulent strains of bacteria: *Salmonella typhosa* and *Pasteurella pestis*. Ann N Y Acad Sci. 1960;88:1125–35 PMID: 13689227

7. Burrows TW, Bacon GA. The basis of virulence in *Pasteurella pestis*: attempts to induce mutation from avirulence to virulence. Br J Exp Pathol. 1954;35(2):129–33 PMID: 13149733

8. Burrows TW, Bacon GA. The effects of loss of different virulence determinants on the virulence and immunogenicity of strains of *Pasteurella pestis*. Br J Exp Pathol. 1958;39(3):278–91 PMID: 13546547

9. Goguen JD, Yother J, Straley SC. Genetic analysis of the low calcium response in *Yersinia pestis* mu d1(Ap lac) insertion mutants. J Bacteriol. 1984;160(3):842–8 PMID: 6094509

10. Yahr TL, Mende-Mueller LM, Friese MB, Frank DW. Identification of type III secreted products of the *Pseudomonas aeruginosa* exoenzyme S regulon. J Bacteriol. 1997;179(22):7165–8. doi: 10.1128/jb.179.22.7165-7168.1997 PMID: 9371466

11. Pettersson J, Holmstrom A, Hill J, Leary S, Frithz-Lindsten E, von Euler-Matell A, et al. The V-antigen of *Yersinia* is surface exposed before target cell contact and involved in virulence protein translocation. Mol Microbiol. 1999;32(5):961–76 PMID: 10361299

12. Sawa T, Ito E, Nguyen VH, Haight M. Anti-PcrV antibody strategies against virulent *Pseudomonas aeruginosa*. Hum Vaccin Immunother. 2014;10(10):2843–52. doi: 10.4161/21645515.2014.971641 PMID: 25483637

13. Baer M, Sawa T, Flynn P, Luehrsen K, Martinez D, Wiener-Kronish JP, et al. An engineered human antibody fab fragment specific for *Pseudomonas aeruginosa* PcrV antigen has potent antibacterial activity. Infect Immun. 2009;77(3):1083–90. doi: 10.1128/IAI.00815-08 PMID: 19103766

14. Faure K, Fujimoto J, Shimabukuro DW, Ajayi T, Shime N, Moriyama K, et al. Effects of monoclonal anti-PcrV antibody on *Pseudomonas aeruginosa*-induced acute lung injury in a rat model. J Immune Based Ther Vaccines. 2003;1(1):2. doi: 10.1186/1476-8518-1-2 PMID: 12943554

15. Frank DW, Vallis A, Wiener-Kronish JP, Roy-Burman A, Spack EG, Mullaney BP, et al. Generation and characterization of a protective monoclonal antibody to *Pseudomonas aeruginosa* PcrV. J Infect Dis. 2002;186(1):64–73. doi: 10.1086/341069 PMID: 12089663

16. Imamura Y, Yanagihara K, Fukuda Y, Kaneko Y, Seki M, Izumikawa K, et al. Effect of anti-PcrV antibody in a murine chronic airway *Pseudomonas aeruginosa* infection model. Eur Respir J. 2007;29(5):965–8. doi: 10.1183/09031936.00147406 PMID: 17301098

17. Katoh H, Yasumoto H, Shimizu M, Hamaoka S, Kinoshita M, Akiyama K, et al. IV Immunoglobulin for acute lung injury and bacteremia in *Pseudomonas aeruginosa* pneumonia. Crit Care Med. 2016;44(1):e12–24. doi: 10.1097/CCM.0000000000001271 PMID: 26317571

18. Kinoshita M, Kato H, Yasumoto H, Shimizu M, Hamaoka S, Naito Y, et al. The prophylactic effects of human IgG derived from sera containing high anti-PcrV titers against pneumonia-causing *Pseudomonas aeruginosa*. Hum Vaccin Immunother. 2016;12(11):2833–46. doi: 10.1080/21645515.2016.1209280 PMID: 27454613

19. Neely AN, Holder IA, Wiener-Kronish JP, Sawa T. Passive anti-PcrV treatment protects burned mice against *Pseudomonas aeruginosa* challenge. Burns. 2005;31(2):153–8. doi: 10.1016/j.burns.2004.09.002 PMID: 15683685

20. Shime N, Sawa T, Fujimoto J, Faure K, Allmond LR, Karaca T, et al. Therapeutic administration of anti-PcrV F(ab’)(2) in sepsis associated with *Pseudomonas aeruginosa*. J Immunol. 2001;167(10):5880–6. doi: 10.4049/jimmunol.167.10.5880 PMID: 11698464

21. Song Y, Baer M, Srinivasan R, Lima J, Yarranton G, Bebbington C, et al. PcrV antibody-antibiotic combination improves survival in *Pseudomonas aeruginosa*-infected mice. Eur J Clin Microbiol Infect Dis. 2012;31(8):1837–45. doi: 10.1007/s10096-011-1509-2 PMID: 22187351

22. Wang Q, Li H, Zhou J, Zhong M, Zhu D, Feng N, et al. PcrV antibody protects multi-drug resistant *Pseudomonas aeruginosa* induced acute lung injury. Respir Physiol Neurobiol. 2014;193:21–8. doi: 10.1016/j.resp.2014.01.001 PMID: 244183531

23. Warrener P, Varkey R, Bonnell JC, DiGiandomenico A, Camara M, Cook K, et al. A novel anti-PcrV antibody providing enhanced protection against *Pseudomonas aeruginosa* in multiple animal infection models. Antimicrob Agents Chemother. 2014;58(8):4384–91. doi: 10.1128/AAC.02643-14 PMID: 24841258

24. Le HN, Quetz JS, Tran VG, Le Vtm, Aguiar-Alves F, Pinheiro MG, et al. MEDI3902 correlates of protection against severe *Pseudomonas aeruginosa* pneumonia in a rabbit acute pneumonia model. Antimicrob Agents Chemother. 2018;62(5). doi: 10.1128/AAC.02565-17 PMID: 29483116

25. Hamaoka S, Naito Y, Katoh H, Shimizu M, Kinoshita M, Akiyama K, et al. Efficacy comparison of adjuvants in PcrV vaccine against *Pseudomonas aeruginosa* pneumonia. Microbiol Immunol. 2017;61(2):64–74. doi: 10.1111/1348-0421.12467 PMID: 28370521

26. Moriyama K, Wiener-Kronish JP, Sawa T. Protective effects of affinity-purified antibody and truncated vaccines against *Pseudomonas aeruginosa* V-antigen in neutropenic mice. Microbiol Immunol. 2009;53(11):587–94. doi: 10.1111/j.1348-0421.2009.00165.x PMID: 19903258

27. Naito Y, Hamaoka S, Kinoshita M, Kainuma A, Shimizu M, Katoh H, et al. The protective effects of nasal PcrV-CpG oligonucleotide vaccination against *Pseudomonas aeruginosa* pneumonia. Microbiol Immunol. 2018;62(12):774–85. doi: 10.1111/1348-0421.12658 PMID: 30378708

28. Ali SO, Yu XQ, Robbie GJ, Wu Y, Shoemaker K, Yu L, et al. Phase 1 study of MEDI3902, an investigational anti-*Pseudomonas aeruginosa* PcrV and Psl bispecific human monoclonal antibody, in healthy adults. Clin Microbiol Infect. 2019;25(5):629 e1–e6. doi: 10.1016/j.cmi.2018.08.004 PMID: 30107283

29. Francois B, Luyt CE, Dugard A, Wolff M, Diehl JL, Jaber S, et al. Safety and pharmacokinetics of an anti-PcrV PEGylated monoclonal antibody fragment in mechanically ventilated patients colonized with *Pseudomonas aeruginosa*: a randomized, double-blind, placebo-controlled trial. Crit Care Med. 2012;40(8):2320–6. doi: 10.1097/CCM.0b013e31825334f6 PMID: 22622405

30. Jain R, Beckett VV, Konstan MW, Accurso FJ, Burns JL, Mayer-Hamblett N, et al. KB001-A, a novel anti-inflammatory, found to be safe and well-tolerated in cystic fibrosis patients infected with *Pseudomonas aeruginosa*. J Cyst Fibros. 2018;17(4):484–91. doi: 10.1016/j.jcf.2017.12.006 PMID: 29292092

31. Milla CE, Chmiel JF, Accurso FJ, VanDevanter DR, Konstan MW, Yarranton G, et al. Anti-PcrV antibody in cystic fibrosis: a novel approach targeting *Pseudomonas aeruginosa* airway infection. Pediatr Pulmonol. 2014;49(7):650–8. doi: 10.1002/ppul.22890 PMID: 24019259

32. Yasumoto H, Katoh H, Kinoshita M, Shimizu M, Hamaoka S, Akiyama K, et al. Epidemiological analysis of serum anti-*Pseudomonas aeruginosa* PcrV titers in adults. Microbiol Immunol. 2016;60(2):114–20. doi: 10.1111/1348-0421.12353 PMID: 26696420

33. Sawa T, Katoh H, Yasumoto H. V-antigen homologs in pathogenic gram-negative bacteria. Microbiol Immunol. 2014;58(5):267–85. doi: 10.1111/1348-0421.12147 PMID: 24641673

34. Liu PV. The roles of various fractions of *Pseudomonas aeruginosa* in its pathogenesis. 3. Identity of the lethal toxins produced in vitro and in vivo. J Infect Dis. 1966;116(4):481–9. Epub 1966/10/01. doi: 10.1093/infdis/116.4.481 PMID: 4959184

35. Sawa T, Corry DB, Gropper MA, Ohara M, Kurahashi K, Wiener-Kronish JP. IL-10 improves lung injury and survival in *Pseudomonas aeruginosa* pneumonia. J Immunol. 1997;159(6):2858–66. Epub 1997/09/23 PMID: 9300709

36. Sawa T, Ohara M, Kurahashi K, Twining SS, Frank DW, Doroques DB, et al. In vitro cellular toxicity predicts *Pseudomonas aeruginos*a virulence in lung infections. Infect Immun. 1998;66(7):3242–9. Epub 1998/06/25 PMID: 9632591

37. Gazi AD, Charova SN, Panopoulos NJ, Kokkinidis M. Coiled-coils in type III secretion systems: structural flexibility, disorder and biological implications. Cell Microbiol. 2009;11(5):719–29. doi: 10.1111/j.1462-5822.2009.01297.x PMID: 19215225

38. Broz P, Mueller CA, Muller SA, Philippsen A, Sorg I, Engel A, et al. Function and molecular architecture of the *Yersinia* injectisome tip complex. Mol Microbiol. 2007;65(5):1311–20. doi: 10.1111/j.1365-2958.2007.05871.x PMID: 17697254

39. Mota LJ. Type III secretion gets an LcrV tip. Trends Microbiol. 2006;14(5):197–200. doi: 10.1016/j.tim.2006.02.010 PMID: 16564172

40. Mueller CA, Broz P, Cornelis GR. The type III secretion system tip complex and translocon. Mol Microbiol. 2008;68(5):1085–95. doi: 10.1111/j.1365-2958.2008.06237.x PMID: 18430138

41. Mueller CA, Broz P, Muller SA, Ringler P, Erne-Brand F, Sorg I, et al. The V-antigen of *Yersinia* forms a distinct structure at the tip of injectisome needles. Science. 2005;310(5748):674–6. doi: 10.1126/science.1118476 PMID: 16254184

42. Sawa T, Yahr TL, Ohara M, Kurahashi K, Gropper MA, Wiener-Kronish JP, et al. Active and passive immunization with the *Pseudomonas* V antigen protects against type III intoxication and lung injury. Nat Med. 1999;5(4):392–8. doi: 10.1038/7391 PMID: 10202927

43. Alonso JM, Vilmer E, Mazigh D, Mollaret HH. Mechanisms of acquired resistance to plague in mice infected by *Yersinia enterocolitica* O:3. Curr Microbiol. 1980;4:117–22

44. Wake A, Maruyama T, Akiyama Y, Yamamoto M. he role of virulence antigen (VW) in the protection of mice against *Yersinia pestis* infection. Curr Microbiol. 1983;8:73–7

45. Roggenkamp A, Geiger AM, Leitritz L, Kessler A, Heesemann J. Passive immunity to infection with *Yersinia spp*. mediated by anti-recombinant V antigen is dependent on polymorphism of V antigen. Infect Immun. 1997;65(2):446–51 PMID: 9009295

46. Motin VL, Nakajima R, Smirnov GB, Brubaker RR. Passive immunity to *yersiniae* mediated by anti-recombinant V antigen and protein A-V antigen fusion peptide. Infect Immun. 1994;62(10):4192–201 PMID: 7927675

47. Moss J, Ehrmantraut ME, Banwart BD, Frank DW, Barbieri JT. Sera from adult patients with cystic fibrosis contain antibodies to *Pseudomonas aeruginosa* type III apparatus. Infect Immun. 2001;69(2):1185–8. doi: 10.1128/IAI.69.2.1185-1188.2001 PMID: 11160019

48. McCarter L. The multiple identities of *Vibrio parahaemolyticus*. J Mol Microbiol Biotechnol. 1999;1(1):51–7 PMID: 10941784

49. Li L, Meng H, Gu D, Li Y, Jia M. Molecular mechanisms of *Vibrio parahaemolyticus* pathogenesis. Microbiol Res. 2019;222:43–51. doi: 10.1016/j.micres.2019.03.003 PMID: 30928029

50. Vandenberghe J, Verdonck L, Robles-Arozarena R, Rivera G, Bolland A, Balladares M, et al. *Vibrios* associated with *Litopenaeus vannamei* larvae, postlarvae, broodstock, and hatchery probionts. Appl Environ Microbiol. 1999;65(6):2592–7 PMID: 10347048

51. Karunasagar I, Pai R, Malathi GR, Karunasagar I. Mass mortality of *Penaeus monodon* larvae due to antibiotic resistant *Vibrio harveyi* infection. Aquaculture 1994; 128:203–9

52. Liu PC, Lee KK, Yii KC, Kou GH, Chen SN. Isolation of *Vibrio harveyi* from diseased kuruma prawns *Penaeus japonicus*. Curr Microbiol. 1996;33(2):129–32 PMID: 8662185

53. Zen-Yoji H, Le Clair RA, Ota K, Montague TS. Comparison of *Vibrio parahaemolyticus* cultures isolated in the United States with those isolated in Japan. J Infect Dis. 1973;127(3):237–41. doi: 10.1093/infdis/127.3.237 PMID: 4569803

54. Zhao Z, Chen C, Hu CQ, Ren CH, Zhao JJ, Zhang LP, et al. The type III secretion system of *Vibrio alginolyticus* induces rapid apoptosis, cell rounding and osmotic lysis of fish cells. Microbiology. 2010;156(Pt 9):2864–72. doi: 10.1099/mic.0.040626-0 PMID: 20576689

55. Zhao Z, Zhang L, Ren C, Zhao J, Chen C, Jiang X, et al. Autophagy is induced by the type III secretion system of *Vibrio alginolyticus* in several mammalian cell lines. Arch Microbiol. 2011;193(1):53–61. doi: 10.1007/s00203-010-0646-9 PMID: 21046072

56. Barillo DJ, McManus AT, Cioffi WG, McManus WF, Kim SH, Pruitt BA, Jr. Aeromonas bacteraemia in burn patients. Burns. 1996;22(1):48–52 PMID: 8719317

57. Juan HJ, Tang RB, Wu TC, Yu KW. Isolation of *Aeromonas hydrophila* in children with diarrhea. J Microbiol Immunol Infect. 2000;33(2):115–7 PMID: 10917882

58. Vila J, Ruiz J, Gallardo F, Vargas M, Soler L, Figueras MJ, et al. *Aeromonas spp*. and traveler’s diarrhea: clinical features and antimicrobial resistance. Emerg Infect Dis. 2003;9(5):552–5 PMID: 12737738

59. Duchaud E, Rusniok C, Frangeul L, Buchrieser C, Givaudan A, Taourit S, et al. The genome sequence of the entomopathogenic bacterium *Photorhabdus luminescens*. Nat Biotechnol. 2003;21(11):1307–13. doi: 10.1038/nbt886 PMID: 14528314

60. ffrench-Constant R, Waterfield N, Daborn P, Joyce S, Bennett H, Au C, et al. Photorhabdus: towards a functional genomic analysis of a symbiont and pathogen. FEMS Microbiol Rev. 2003;26(5):433–56. doi: 10.1111/j.1574-6976.2003.tb00625.x PMID: 12586390

61. Silva CP, Waterfield NR, Daborn PJ, Dean P, Chilver T, Au CP, et al. Bacterial infection of a model insect: *Photorhabdus luminescens* and *Manduca sexta*. Cell Microbiol. 2002;4(6):329–39 PMID: 12067318

62. Waterfield NR, Daborn PJ, ffrench-Constant RH. Genomic islands in *Photorhabdus*. Trends Microbiol. 2002;10(12):541–5 PMID: 12564983

63. Gerrard JG, McNevin S, Alfredson D, Forgan-Smith R, Fraser N. *Photorhabdus* species: bioluminescent bacteria as emerging human pathogens? Emerg Infect Dis. 2003;9(2):251–4. doi: 10.3201/eid0902.020222 PMID: 12603999

